# Synchronization of two active ears via binaural coupling

**DOI:** 10.1101/2025.11.20.689396

**Authors:** Rebecca E. Whiley, Filipe Ledo, Olha Fedoryk, Christopher Bergevin

**Affiliations:** Department of Biology, York University, Toronto, ON, Canada; Department of Medical Biophysics, University of Western Ontario, London, ON, Canada; Department of Physics & Astronomy, York University, Toronto, ON, Canada

**Keywords:** otoacoustic emissions, synchrony, active hearing, sound localization

## Abstract

The two ears of many non-mammalian vertebrates are acoustically coupled through an interaural cavity. Sounds can hit both the external and internal surfaces of the tympanic membranes, providing mechanical directional sensitivity for sound localization. This coupling may also give rise to binaural synchronization. In other words, each inner ear could influence the function of the other. Previous work in lizards demonstrates that binaural coupling affects the spontaneous otoacoustic emissions (SOAEs) measured at each external auditory meatus, with properties indicative of binaural synchronization. However, it is unclear how binaural coupling and, consequently, synchronization contribute to SOAE generation, which is typically modelled as being localized to an individual ear. We simultaneously measured SOAEs at both ears of green anole lizards (*Anolis carolinensis*) and found robust relationships between them, including evidence that binaural synchronization could not be attributed to one ear uniformly driving the other. Instead, we observed frequency-dependent phase-locking between the two ears, primarily at frequencies where SOAE peaks occurred in both ears. While some pairs of ears were more strongly synchronized than others, we consistently found that binaural coupling could have greater effects on the resulting emissions than the coupling between generators within the same ear. We propose a framework for active hearing that incorporates binaural coupling, accounting for its effects on SOAE generation and sound localization in the green anole.

## 1. Introduction

Bilateral symmetry has important implications for sensory perception. In many vertebrates, comparisons between sounds at the left and right ears create interaural cues that contribute to binaural sound localization (Fay and Popper, 2005). While interaural cues are processed neurally in mammals (Grothe et al., 2010), they can be processed at the auditory periphery in some non-mammals (Dooling et al., 2000). In lizards, there is a direct acoustic pathway between each ear’s tympanic membrane (TyM), forming an interaural cavity (IAC) where mechanical coupling can occur (see schematic in Figure 1A; Christensen-Dalsgaard, 2005). Each TyM receives acoustic cues from outside and inside the head, and this binaural coupling forms a “pressure-difference receiver” system (Christensen-Dalsgaard and Manley, 2008; Christensen-Dalsgaard et al., 2021; Fletcher and Thwaites, 1979). As a result, peripheral comparisons between sounds can serve as the basis of binaural sound localization prior to auditory transduction in the inner ear.

**Figure 1:**
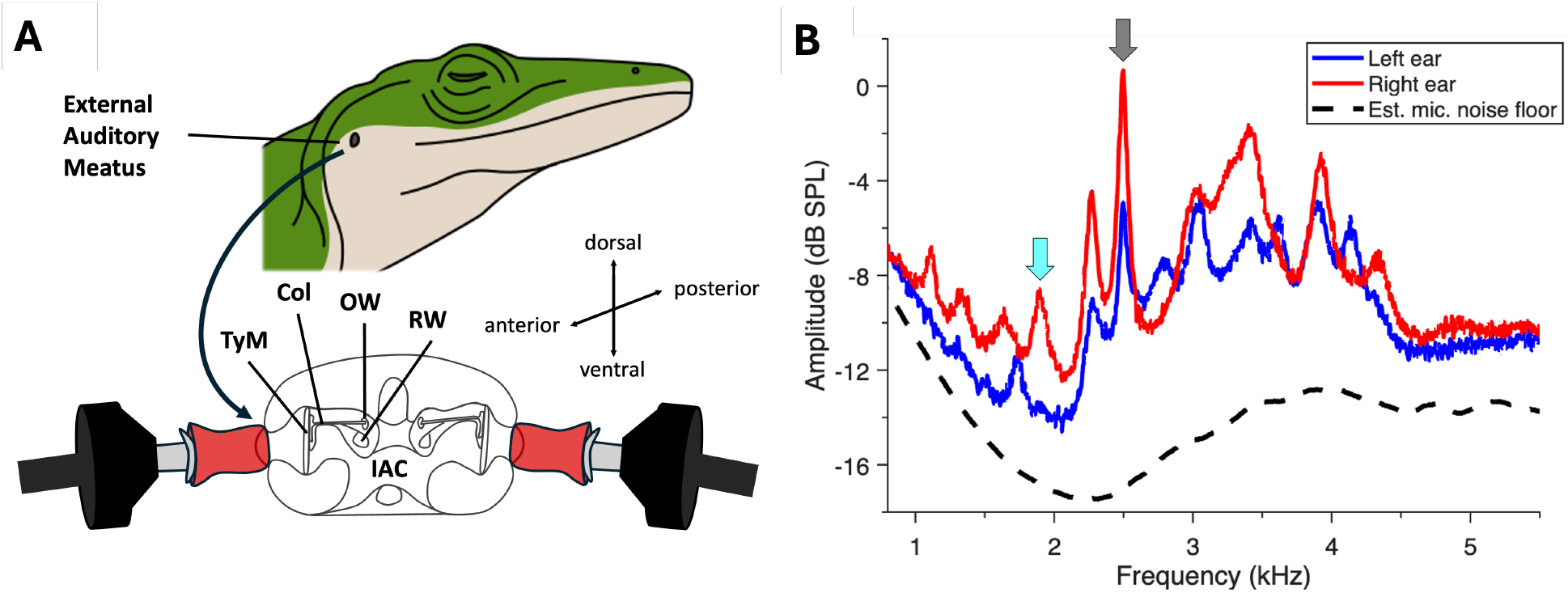
(**A**) Schematic of simultaneous binaural SOAE measurement paradigm in a green anole lizard using an ER-10C probe coupled to each external auditory meatus (top) and a coronal cross-section of the head (bottom). **IAC**: interaural cavity; **TyM**: tympanic membrane; **Col**: columella; **RW**: round window; **OW**: oval window. The inner ear is found behind the oval window. Anatomical diagram adapted from Wever (1978). (**B**) Simultaneously measured bilateral spontaneous otoacoustic emissions (SOAEs) from the green anole lizard’s left (blue) and right (red) ears. The estimated microphone noise floor is illustrated by the black dashed line. The grey arrow at 2.5 kHz illustrates a case where peaks occurred at the same frequency in both ears (bilateral matched peak) while the cyan arrow at 1.9 kHz illustrates a case where a peak occurred at that frequency only in one ear (unmatched peak).

Binaural coupling could similarly drive relationships between the active mechanics of the two inner ears. Within the inner ear, active force generation is utilized to improve functionality: metabolic energy is used to enhance the ear’s sensitivity and frequency selectivity to external sounds (Hudspeth, 2008). As a byproduct of this active process, acoustic energy exits the inner ear and induces oscillations of the TyM, producing sounds that are measurable in the ear canal (Kemp, 1979). Such oscillations can occur in the absence of external stimulation, resulting in subthreshold sounds called spontaneous otoacoustic emissions (SOAEs; Bergevin et al., 2017). SOAEs commonly manifest as arrays of spectral peaks that are unique to an individual ear and are observed across vertebrates, including humans and non-mammals like lizards (Bergevin et al., 2015; Manley, 2000). Given that each inner ear is an active, self-oscillating system, binaural coupling could lead to binaural synchronization. In other words, the ears could synchronize because of their weak interaction via the IAC, causing the oscillators of each inner ear to adjust their characteristic frequencies (Pikovsky et al., 2001). Consequently, we might predict that binaural synchronization contributes to SOAE generation, and an emission measured at one ear is the product of both.

Several recent studies in lizards indicate that an emission could derive from two ears due to binaural coupling. Direct measurements of SOAE activity inside the IAC of green anole lizards (*Anolis carolinensis*) support its role in binaural coupling (Bergevin et al., 2020). SOAEs measured by placing a probe microphone in the mouth (presumably capturing contributions from both ears) were compared to those measured at each meatus, revealing peaks at similar frequencies on both sides of the TyM. Acoustic modelling based on IAC morphology, characterized by µCT (Gauthier et al., 2012), showed that the motion of one TyM, presumably driven by the ipsilateral inner ear, was sufficient to drive the contralateral TyM and thereby couple the two inner ears (Bergevin et al., 2020). When simultaneously measuring responses at each external auditory meatus in tokay geckos (*Gekko gecko*), Roongthumskul et al. (2019) observed that SOAE peaks at similar frequencies in both ears often coincided with regions of synchronization between the two waveforms. By manipulating binaural coupling, they demonstrated that some SOAE peaks arise specifically due to binaural synchronization; these peaks were abolished by suppressing a peak at the same frequency in the contralateral ear.

Binaural synchronization would imply that the emissions measured at each external auditory meatus are determined by the properties of both active ears and their coupling via the IAC. We evaluated whether binaural synchronization affects SOAE generation in green anole lizards by simultaneously measuring SOAEs at each external auditory meatus. We extended the vector strength analysis methods previously applied to geckos (Roongthumskul et al., 2019) and incorporated fluctuation analyses (Bergevin et al., 2025) as a secondary method for investigating binaural synchronization at frequency bands corresponding to filtered SOAE peaks. Together, our analysis techniques allowed us not only to characterize the strength of the relationships between ears but also the temporal causality (e.g., delays), which could be used to infer whether binaural synchronization occurs because one ear drives the other. Based on our results, we propose that models of the anole’s auditory system should account for the possibility that there could be two active ears, not just one, contributing to the data.

## 2. Methods

Experiments were conducted on eight adult male green anole lizards (*Anolis carolinensis*). Each lizard was anesthetized with an injection of sodium pentobarbital (Nembutal or Euthanyl at 30–36 mg/kg, *N* = 5) or alfaxalone (Alfaxan at 27.5 mg/kg, *N* = 3) and placed in an acoustic isolation booth. Body temperature was monitored and, if necessary, a regulated heating pad was used to stabilize temperature. All procedures were approved by the Animal Care Committee at York University.

### 2.1. Calibration and SOAE Measurements

Measurements were made with a custom data acquisition system using a 24-bit sound card (C++ on Lynx TWO-A, Lynx Studio Technology, or LabVIEW on NI PXI-4461, National Instruments). We verified that these systems were functionally equivalent by comparing responses from the same subjects across the two sound cards. Signals were output and recorded at a sampling rate of 44.1 kHz using two ER-10C probes (Etymotic Research), with the microphone output amplified by 40 dB and high-pass filtered at 410 Hz. Each ER-10C probe was close-coupled to the lizard’s external auditory meatus using grease (Figure 1A). Probe earphones were calibrated *in situ* with random-phase flat-spectrum noise. We averaged the responses across 10 presentations and computed the transfer function to screen for amplitude drop-offs below 2 kHz, indicative of a gap in the probe’s coupling. When necessary, we re-coupled the probe and repeated the calibration until the gap was resolved. For half of the lizards, we performed trans-IAC calibrations (presenting noise to one ear and recording the response from the contralateral ear) and referenced the transfer functions to the ipsilateral transfer functions to characterize interaural sound transmission at the external surfaces of the TyMs.

SOAE waveforms were acquired simultaneously from both ears for a duration of 120 seconds and, unless otherwise specified, analyzed offline in MATLAB version 2025a. SOAE peaks were identified from spectrally averaged amplitudes (400 segments of 8192 samples, frequency resolution of 5.4 Hz) and isolated with a recursive exponential filter in the spectral domain (Bergevin et al., 2025; Shera and Zweig, 1993). The filter’s bandwidth set the boundaries of each peak; it was centered to capture the elevated spectrum around a local maximum before it entered plateaus at local minima without overlapping the filters for neighboring peaks. Consequently, the peak’s center frequency was defined by the filter and could be slightly offset from the peak’s local maximum. For simplicity, SOAE “valleys” were defined as the plateau regions between the filters of adjacent peaks.

We used the filtered peak’s signal-to-noise ratio (SNR) as a proxy for estimating SOAE peak amplitude. The SNR was calculated as the difference between the maximum amplitude and the average amplitude at the filter flanks (i.e., the center frequency 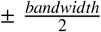. Because we calculated the SNR as the peak maximum relative to the adjacent valleys, which often sit above the estimated microphone noise floor, the SNR values are likely underestimates of SOAE amplitude (note how the valleys in the SOAE spectra [blue and red lines] sit above the noise floor [grey dashed line] in Figure 1B). SOAE peaks with SNRs of at least 1.5 dB were considered for further analysis. While this was a more generous inclusion criterion than commonly used for SOAE peak identification, it was chosen to maximize comparisons between waveforms.

We defined “bilateral matched peaks” as SOAE peaks for which the same filter could be used to encapsulate the peaks in both ears (see grey arrow in Figure 1B). To facilitate further comparisons between bilateral waveforms, SOAE peaks with SNRs between 1–1.5 dB were included if there was a match ≥1.5 dB in the contralateral ear’s SOAE spectrum. If an SOAE peak did not have a match in the contralateral ear and its SNR was ≥ 1.5 dB, it was considered an “unmatched peak” (see cyan arrow in Figure 1B). We also evaluated “bilateral matched valleys”, representing regions where there were valleys between bilateral matched peaks in both ears or an unmatched peak in one ear coincided with a valley in the contralateral ear.

### 2.2. Vector Strength Analyses

#### 2.2.1. Vector Strength

Vector strength was used to measure the entrainment between bilateral SOAE waveforms on a scale of 0 to 1, with a value of 1 indicating perfect binaural synchronization (Roongthumskul et al., 2019). Both SOAE waveforms were high-pass filtered with a second-order Butterworth filter at 200 Hz to eliminate residual line noise artifacts. We calculated the phase differences between waveforms at each frequency *f* as the argument of the ratio of the left ear’s complex spectrum (*F*_*L*_) to the right ear’s complex spectrum (*F*_*R*_) in segments (denoted *j*) of 8192 samples, obtaining

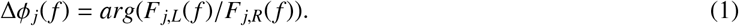

We computed the vector strength, *V*, by taking the absolute value of the argument of the ensemble average of Δ*ϕ*_*j*_( *f* ) over 645 pairs of non-overlapping segments as

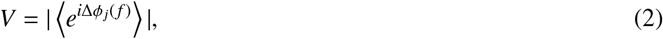

where *V* ∈ [0, 1]. The phase differences of the mean Δ*ϕ*( *f* ) were calculated as the argument of 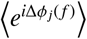, where Δ*ϕ* ∈ [− 0.5, 0.5] cycles. We applied a 1-cycle (i.e., 2π) adjustment to values below 0 such that Δ*ϕ* ∈ [0, 1].

For independent signals, vector strength would be near 0, and phase differences (Δ*ϕ*) would be variable with time. We verified that asynchronously measured binaural waveforms had noisy Δ*ϕ* and vector strength values indistinguishable from the noise (all *V* < 0.20). Vector strength was considered elevated and indicative of binaural synchronization when *V* ≥ 0.30. We defined the vector strength and average phase difference for an SOAE peak as the maximum value of *V* and mean value of Δ*ϕ* when *V* ≥ 0.30 within ±15 Hz of the peak’s center frequency.

#### 2.2.2. Time-Delayed Vector Strength

We considered how binaural synchronization varied as a function of a temporal offset, termed *ξ* (ms), between the two ears by calculating the time-shifted vector strength *V*_*ξ*_ (computed in Python version 6.1). Waveforms were high-pass filtered and parsed into segments of 8192 samples as described in Section 2.2.1, and then we applied a Gaussian filter (full width at half maximum of 471 samples) to each segment to reduce temporal overlap between waveforms. Rather than comparing the phase differences between waveforms at identical points in time, we adapt Equation 1 such that we take the difference between the angle of the left ear’s complex spectrum and the angle of the right ear’s time-shifted complex spectrum. Our phase difference is now a function of *f* and time *t*, 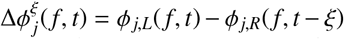, with the offset between segments defined by *ξ*. This offset could be positive or negative, but *ξ* was always less than the duration of *t*. Positive values of *ξ* represent the left ear’s complex spectrum leading the right, while negative values represent the converse, and higher values of *ξ* indicate greater temporal offset between the segments of each waveform. *V*(*ξ*) was subsequently computed by subbing 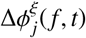 into Equation 2 across all *ξ*. To determine the value of *ξ* that produced the greatest vector strength, we fit Gaussian functions to *V*(*ξ*) at frequency bands at bilateral matched peaks. We designated the time-shifted vector strength *V*_*ξ*_ as the maximum value of the fitted function (with the same inclusion criterion as *V*: 0.30), and *ξ* at the center of the maximum was the *V*_*ξ*_ delay.

#### 2.2.3. Time-Delayed Phase Coherence

We adapted the computation of vector strength to measure self-coherence within a single SOAE waveform (Peacock et al., 2025). We referenced the SOAE signal to itself at a previous instance in time (specified by the temporal offset *ξ*), obtaining the time-delayed phase coherence *C*_*ξ*_ as a measure of SOAE phase stability over time (computed in Python version 6.1). After waveforms were high-pass filtered, they were parsed into segments of 2048 samples (frequency resolution of 21.5 Hz) and Gaussian filtered (full width at half maximum of 471 samples). While shorter segments resulted in a lower frequency resolution than *V* and *V*_*ξ*_, longer segments were undesirable because phase coherence decays over time due to the time-frequency tradeoff inherent to the Fourier transform (Peacock et al., 2025). Similar to our computation of *V*_*ξ*_, we adapted Equation 1 to take the difference of the angle of one ear’s complex spectrum to the angle of the same ear’s time-shifted complex spectrum. Here, phase differences are still a function of *f* and time *t*, but are calculated within one waveform: 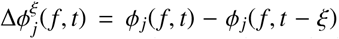. Unlike *V*_*ξ*_, the offset between segments *ξ* was fixed at 17.4 ms (equivalent to 768 samples). The time-delayed phase coherence *C*_*ξ*_ was obtained by subbing the within-ear phase differences 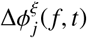 into Equation 2. The criterion for defining *C*_*ξ*_ “peaks” was values ≥0.05, which was more generous than for vector strength due to the noisier nature of computing the phase relationships within waveforms.

### 2.3. Filtered Peak Fluctuation Correlation Analyses

We evaluated SOAE temporal fluctuations following the methodology of Bergevin et al. (2025) and refer the reader there for a more detailed description of the analysis paradigm. While the vector strength analyses described in Section 2.2 were agnostic to the presence of SOAE peaks, the methods here explicitly depend on the identification and filtering of SOAE peaks and valleys as described in Section 2.1. First, we used the Hilbert transform to compute the analytic signal representation of the filtered peaks or valleys. The amplitude modulation (AM) and frequency modulation (FM) waveforms were obtained from the analytic signal’s envelope and instantaneous frequency, which was computed as the temporal gradient of the signal’s unwrapped phase, respectively.

Correlations between two signals’ AM or FM fluctuations (i.e., AM_1_⋆AM_2_ or FM_1_⋆FM_2_) were computed in the time domain. We took segments *j* with length ℓ of 4096 samples of each signal’s AM or FM and subtracted out the mean value of the signal in each segment to obtain the (real-valued) residual fluctuations Φ _*j*_. The time-domain cross-covariance *T* between the residual fluctuations for two signals was computed for each segment as a function of the lag between samples *τ* as *T*_*j*_(*τ*) = ∑ _*n*_ Φ_1 *j*_(*t* + *τ*)Φ_2 *j*_(*t*). Here, *n* is the total number of overlapping samples (∼ 2ℓ − 1) comprising the segments, and *t* represents time. We normalized *T*_*j*_ using the autocorrelations (e.g., *A*_1 *j*_(*τ*) = ∑ Φ_1 *j*_(*t* + *τ*)Φ_1 *j*_(*t*)) at zero lag to obtain the cross correlation at each segment as 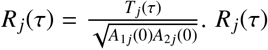 was averaged across segments to obtain the mean normalized cross correlation *R*(*τ*), where *R* ∈ [−1, 1]. Negative functions indicated that when one fluctuation increased, the other decreased, whereas positive functions indicated that both varied in a similar fashion.

We evaluated AM and FM fluctuation correlations for pairs of SOAE peaks and valleys within and across waveforms. Representative fluctuation correlation functions for one subject are illustrated in Figure 2B–D for *R*_*AM*_(*τ*) (top) and *R*_*FM*_(*τ*) (bottom). Within an SOAE waveform, we evaluated pairs of ipsilateral adjacent peaks that represented spectrally adjacent peaks (i.e., nearest neighbors), including those with and without matches in the contralateral ear (Figure 2B). Ipsilateral correlations were computed as the lower-frequency peak relative to the higher-frequency peak, such that positive correlation delays represent the lower frequency leading the higher. Between SOAE waveforms, we evaluated bilateral matched peaks (Figure 2C) and bilateral matched valleys, including frequency bands where valleys occurred in both ears and bands in which unmatched peaks coincided with valleys in the contralateral ear (Figure 2D). Bilateral correlations were computed as the left ear relative to the right ear, such that positive correlation delays represent the left ear leading the right.

**Figure 2:**
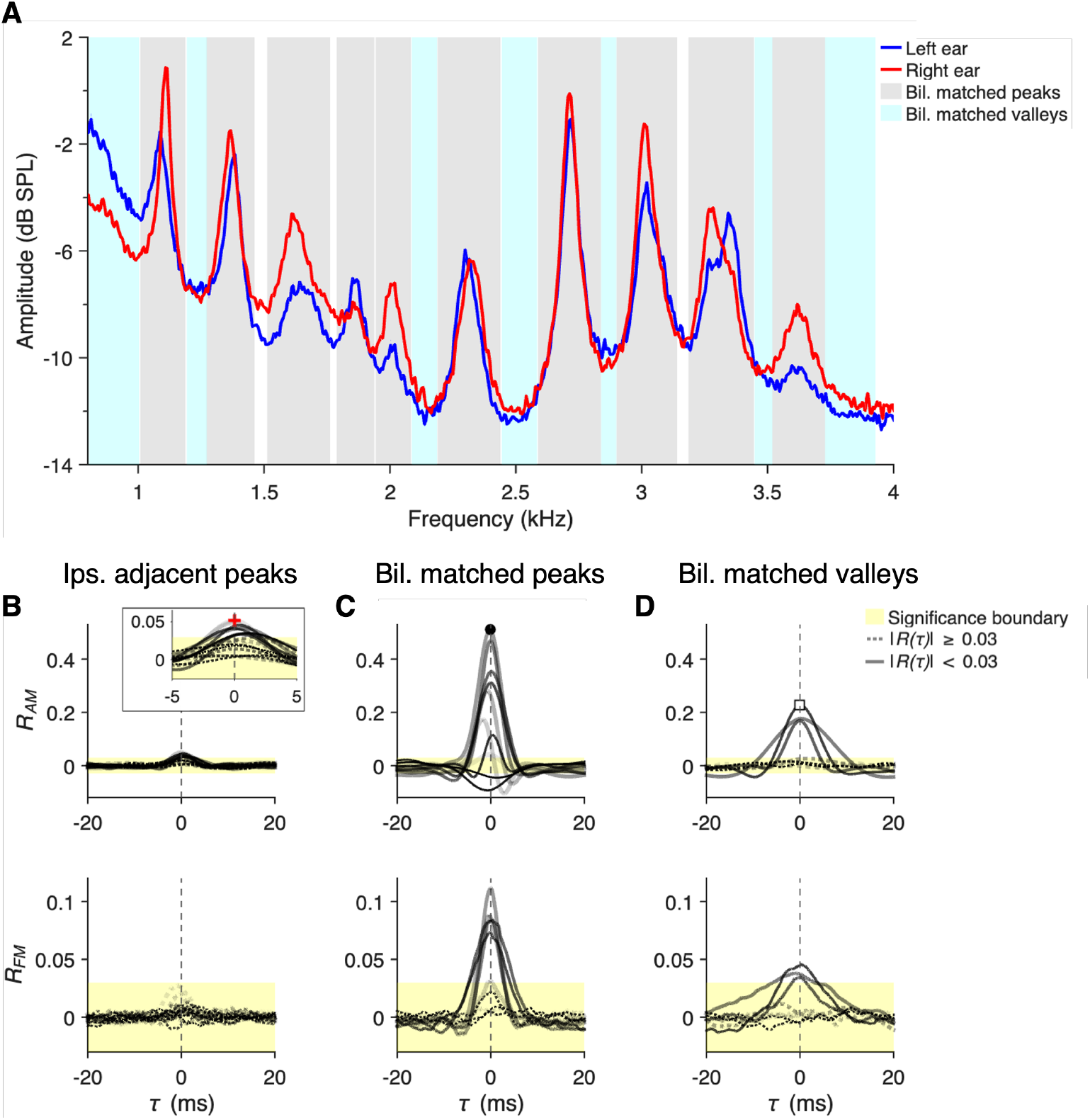
SOAE peak envelope (AM) and instantaneous frequency (FM) fluctuation correlations were evaluated within and between ears. (**A**) SOAE spectra measured at the left (blue) and right (red) ears of a representative green anole lizard. Grey and cyan shading defines the boundaries of the filters for bilateral matched peaks and valleys, respectively. Ipsilateral adjacent peak pairs were obtained from the adjacent grey shaded regions for the left and right ears. (**B–D**) Fluctuation correlation functions between SOAE peak AM (*R*_*AM*_, top) and FM (*R*_*FM*_, bottom) waveforms for pairs of ipsilateral adjacent peaks (**B**), bilateral matched peaks (**C**), and bilateral matched valleys (**D**) for the same subject in panel (**A**). The inset in the top panel of (**B**) provides a better illustration of the correlations between ipsilateral adjacent peaks. Correlation functions with maximum values ≥ ± 0.03 (indicated by yellow shading) were plotted as solid lines, while sub-threshold correlations were plotted as dotted lines. Lines become thicker and less opaque as peak frequency increases. The maximum value of each correlation function was designated the correlation strength and the corresponding *τ* value was the delay. The markers in the top panels of (**B–D**) illustrate representative correlation strength values (red cross, black circle, and white square, respectively). The vertical dashed lines at *τ* = 0 ms identify the boundary between positive delays, indicative of the lower frequency peak leading the higher frequency peak in panel (**B**) and the left ear leading the right in panels (**C–D**), and negative delays, vice versa.

The maximum values of the |*R*(*τ*)| functions were designated as the “AM correlation strength” or “FM correlation strength”. The corresponding *τ* values at the maxima of *R* were designated as the correlation delays (ms). Fluctuation correlations were considered present if the correlation strength was ≥ |0.03|. This threshold criterion was determined by bootstrapping the correlation (see Bergevin et al., 2025). If the correlation function *R*(*τ*) had a single well-defined maximum or minimum, the relationship was classified as positive or negative as per the sign of the correlation strength. Otherwise, a correlation was classified as “indeterminate” to reflect the more complex relationship between fluctuations.

## 3. Results

### 3.1. Interaural Sound Transmission

We characterized interaural sound transmission by comparing transfer functions obtained in response to ipsilateral and contralateral sound stimuli. Our calculations indicated frequency-dependent attenuation of approximately 13.1– 27.2 dB SPL (mean −17.7 ± 4.3 dB SPL) between 1.0–5.0 kHz (top row of Figure A.8), which falls within the range of best sensitivity for the green anole lizard (Brittan-Powell et al., 2010). The corresponding unwrapped phase values decreased with increasing frequency (middle row of Figure A.8). . We estimated passive sound propagation delays to be approximately 0.09 ms (see dashed lines in the bottom panels of Figure A.8) based on the two probe microphones being approximately 3 cm apart: 1 cm between the two TyMs (i.e., the width of the IAC), and 1 cm between each TyM and microphone (see schematic in Figure 1A). At this distance, the delays attributable to sound propagation alone were shorter than the phase-gradients, which ranged from approximately -0.096–0.38 ms (mean 0.141 ± 0.035 ms) at most frequencies between 1.0–5.0 kHz (bottom row of Figure A.8).

### 3.2. Relationships Between Bilateral SOAE Waveforms

SOAE spectra obtained from simultaneous binaural recordings in eight green anole lizards revealed a series of peaks and valleys unique to each subject (see spectra from subject 1 in Figure 3A and all subjects in Figure A.9). Although peaks also occur in the SOAE phase (see Peacock et al., 2025), unless otherwise specified, discussions of “SOAE peaks” and their associated properties refer to the 134 peaks observed in the amplitudes of the Fourier transforms. Spectra from each ear exhibited similarities within a given subject, with each having SOAE peaks at the same frequencies in both ears (i.e., bilateral matched peaks; see grey shading in Figure 3A). Peak center frequencies ranged from 0.9–4.4 kHz (mean 2.5 ± 0.9 kHz) and SNR ranged from 1.1–11.1 dB (mean 4.4 ± 2.5 dB), with higher values occurring near the mean center frequency (Figure 3B).

**Figure 3:**
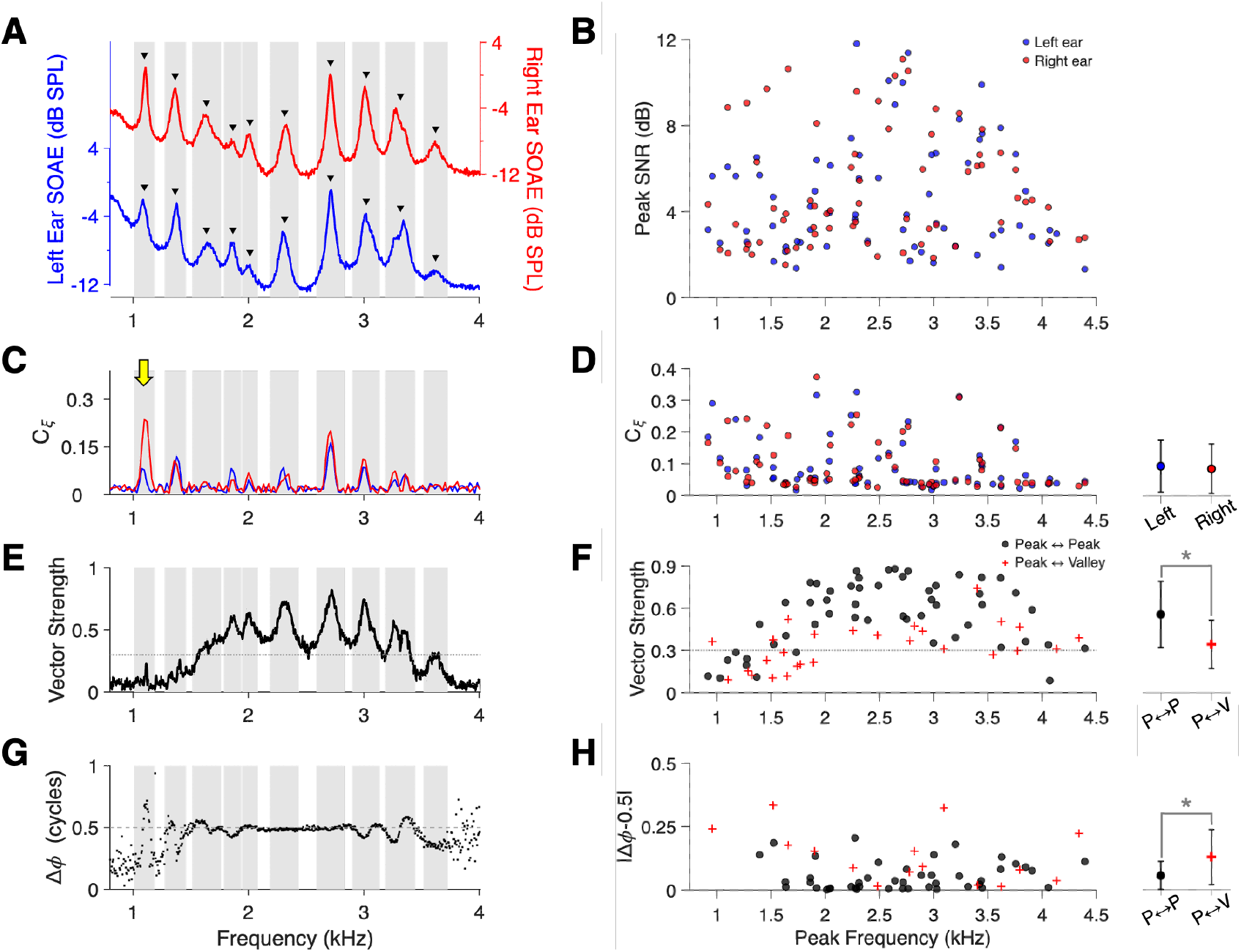
Evaluation of narrowband SOAE properties in a representative green anole lizard (subject 1, panels A, C, E, and G) and all individuals (panels B, D, F, and H). Properties in all subjects are illustrated in Figure A.9 **(A)** SOAE spectra measured simultaneously from the left (blue) and right (red) ears were averaged with a frequency resolution of 5.4 Hz. Note that the different vertical axes for each ear were offset to avoid visual overlap. SOAE peaks are identified by black triangles above each waveform. When SOAE peaks were filtered at the same frequencies in both ears, they were classified as bilateral matched peaks and grey shading defines their bandwidths across all panels. All SOAE peaks illustrated here were classified as bilateral matched peaks. **(B)** SOAE peak signal-to-noise ratios (SNR; dB) across peak frequency (kHz) from the left (blue) and right (red) ears. **(C)** The time-delayed phase coherence (*C*_*ξ*_) for the left (blue) and right (red) ears for subject 1. The yellow arrow at 1.1 kHz highlights the differences in *C*_*ξ*_ peaks between ears. **(D)** *C*_*ξ*_ across peak frequencies from the left (blue) and right (red) ears. Mean values for each ear are plotted with standard deviations (whiskers) in the right panel. There was no significant difference between ears as per a paired *t*-test. **(E)** Vector strength was computed as the absolute value of the averaged phase differences between the simultaneously measured SOAE waveforms. The dashed grey line at 0.30 represents the boundary for identifying peaks in vector strength. **(F)** Vector strength across SOAE peak frequencies for all subjects for bilateral matched peaks (Peak ↔ Peak, black circles) and peaks without matches in the contralateral ear (“unmatched” peaks; Peak ↔ Valley, red crosses). Mean values with standard deviations are plotted in the right panel, with the bracket and asterisk identifying significant differences as per an unpaired *t*-test. Vector strengths were significantly higher in bilateral matched peaks. **(G**) The phase differences (Δ*ϕ*) between binaural SOAE waveforms were approximately ±0.5 cycles when vector strength exceeded 0.30 in subject 1. At low vector strengths, phase differences were variable and closer to 0 or, equivalently, 1 cycle. **(H)** Δ*ϕ* was referenced to 0.5 cycles, such that values closer to 0 indicate phase differences closer to 0.5 cycles. Only phase differences where the corresponding vector strengths were ≥ 0.3 were included. The relative phase differences were closer to 0 for most pairs of bilateral matched peaks (black circles) and closer to 0.5 for unmatched peaks (red crosses), particularly below 2 kHz. Mean values with standard deviations are plotted in the right panel, with the bracket and asterisk identifying significant differences as per an unpaired *t*-test. Phase differences were significantly closer to 0.5 cycles in bilateral matched peaks.

The time-delayed phase coherence (*C*_*ξ*_) revealed peaks at the same frequencies as many SOAE peaks (Figure 3C), particularly at those that were bilaterally matched (grey shading). While *C*_*ξ*_ values at the center frequencies of bilateral matched peaks were similar between ears (paired *t*-test, *t*_(105)_ = 0.104, *p* = 0.917; Figure 3D), differences were apparent at some bilateral matched peaks when comparing data from left (blue) and right (red) ears (Figure 3C). For example, see the yellow arrow at 1.1 kHz in Figure 3C where the *C*_*ξ*_ peak from the right ear was higher in amplitude and frequency than that from the left ear (additional cases are also highlighted by yellow arrows in Figure A.9). Furthermore, *C*_*ξ*_ peaks were absent for many SOAE peaks (see Figure A.9), particularly if a peak had low a SNR or a broad BW. Consistent with this observation, *C*_*ξ*_ was greater in narrower SOAE peaks (inverse relationship with BW, *r*_(132)_ = − 0.34, *p* < 0.001) and those with higher SNR values (*r*_(132)_ = 0.60, *p* < 0.001).

Vector strength peaks typically coincided with the frequencies of bilateral matched peaks (see grey shading in Figure 3E; all subjects in Figure A.9). However, vector strengths could be low even when bilateral matched peak pairs occurred, as shown for the two pairs below 1.5 kHz in Figure 3E. Conversely, vector strength could exceed threshold for unmatched SOAE peaks or in absence of SOAE peaks altogether, particularly at the valleys between bilateral matched peaks (Figure A.9). Most SOAE peaks were part of a bilateral matched peak pair (*N* = 53 pairs), but five subjects had unmatched peaks (*N* = 28) which were the basis of ‘peak–valley’ evaluations of vector strengths (red crosses; Figure 3F). Relative to bilateral matched peaks (i.e., ‘peak–peak’ comparisons, black circles), vector strength was significantly lower for unmatched peaks (unpaired *t*-test assuming equal variance, *t*_(79)_ = 4.24, *p* < 0.001; right panel in Figure 3F). Vector strength was generally elevated in the range of SOAE peaks, being highest near the mean center frequency (Figure 3F). Similar to the relationship between SOAE peak frequency and SNR (Figure 3B), vector strength increased in association with peak SNR for bilateral matched peaks (*r*_(51)_ = 0.574, *p* < 0.001), but no correlation occurred for unmatched peaks.

As shown in Figure 3G, phase differences were typically flat and close to 0.5 cycles when vector strength exceeded threshold (see dashed line at 0.30 in panel E). This was observed across subjects (see Figure A.9), indicative of anti-phase relationships between binaural SOAE waveforms in these regions. When vector strength was below threshold, phase differences were more difficult to interpret due to their high variability (e.g., see values below 1.5 kHz in Figure 3G). We referenced the phase differences to 0.5 cycles (i.e., |Δ*ϕ* − 0.5|), including only peaks for which the vector strength met the threshold criterion. Relative phase differences were frequency-independent for both categories, but were significantly closer to 0 for bilateral matched peaks than for unmatched peaks (unpaired *t*-test assuming unequal variance, *t*_(18.1)_ = − 2.58, *p* = 0.019; right panel in Figure 3H).

We computed the time-delayed vector strength to evaluate binaural synchronization as a function of the delay between the SOAE waveforms measured at each ear. In Figure 4A, the vector strength is plotted for a representative subject as a function of the time delay *ξ* and frequency, with color indicative of strength intensity. As expected, vector strength was higher near the frequencies of bilateral matched peaks (triangle markers in Figure 4A). However, the maximum values at each SOAE peak, *V*_*ξ*_, did not necessarily occur at *ξ* = 0 ms (see square markers). Sometimes vector strength was highest when the two SOAE waveforms were offset in time, consistent with one ear leading the other. When the *V*_*ξ*_ delays were plotted across SOAE peak frequencies (Figure 4B), we observed that delays for a given subject could be positive or negative at different frequency bands. Most *V*_*ξ*_ delays greater than ±1 ms occurred below 1.5 kHz where *V*_*ξ*_ was often ill-defined, with vector strengths below threshold (see values with crosses through them in Figure 4B).

**Figure 4:**
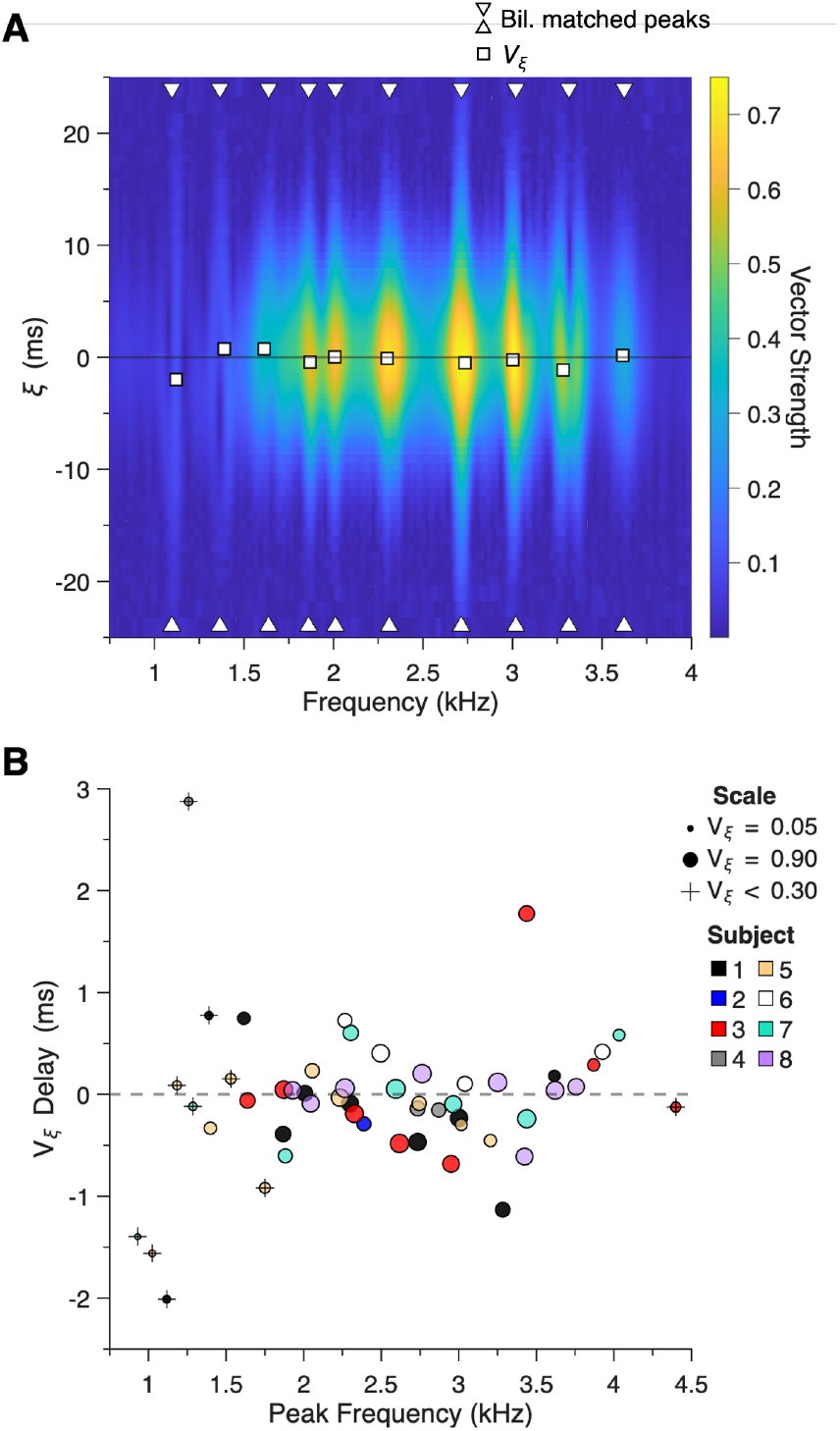
Vector strength between simultaneously measured SOAE waveforms could be greatest when one waveform led the other. (**A**) Illustration of vector strength as a function of the time delay between waveforms, *ξ* (ms), and frequency (kHz) for a green anole lizard (subject 1, black markers in panel **B**). The color bar to the right indicates the intensity of the vector strength. For a given pair of bilateral matched peaks (white triangles at the boundaries of the ordinate axis), the maximum vector strength as a function of *ξ, V*_*ξ*_ (white squares), could occur when there was a temporal offset between waveforms such that one led the other. (**B**) *V*_*ξ*_ delays relative to bilateral matched peak frequencies for all subjects (see legend). Marker size was scaled according to the value of *V*_*ξ*_, with the range specified in the legend. Delays where the corresponding *V*_*ξ*_ was less than the threshold value of 0.3 are identified with a cross through them. The grey dashed horizontal line represents 0 ms delay.

### 3.3. Relationships Between Bilateral SOAE Peaks

As summarized in Table 1, we computed correlations between fluctuations in SOAE peak envelopes (amplitude modulation, AM) or instantaneous frequencies (frequency modulation, FM) to characterize their interactions within (ipsilateral adjacent peak pairs) and between ears (bilateral matched peak or valley pairs). Across pair categories, AM correlations were more common and stronger than FM correlations, which only occurred in bilateral matched pairs (i.e., none were observed between ipsilateral adjacent peak pairs) and when the pair also had a significant AM correlation (Table 1). Significant correlations were most common between bilateral matched peaks, followed by bilateral matched valleys, with the fewest correlations observed between ipsilateral adjacent peaks.

**Table 1:** Summary of SOAE fluctuation correlations between signal envelopes (amplitude modulation, AM) and instantaneous frequencies (frequency modulation, FM) for eight green anole lizards. Correlation strength represents the maximum value of the function *R*(*τ*), where *τ* is the lag between signals. Positive correlation strength is indicative of cases where an increase or decrease in the fluctuation in one signal is accompanied by the same change in the other. Correlation delay (ms) represents the absolute value of the lag at the correlation strength. Positive delays represent the proportion of significant correlation strengths where *τ*≥ 0 ms, indicative of the lower frequency signal leading the higher frequency signal for ipsilateral pairs or the left ear’s signal leading the right ear’s signal for bilateral pairs. Strengths and delays for significant correlations are presented as mean values with standard deviations.

| Category | N | Fluctuation Type | Proportion Significant ($\geq \pm 0.03$) | Proportion Positive ($\geq 0.03$) | Correlation Strength | Proportion Positive Delays ($\geq 0$ ms) | Absolute Correlation Delay (ms) |
| --- | --- | --- | --- | --- | --- | --- | --- |
| Ipsilateral adjacent peaks | 122 | AM | 28% | 97% | $0.050 \pm 0.023$ | 97% | $0.582 \pm 0.532$ |
|  |  | FM | 0% | - | - | - | - |
| Bilateral matched peaks | 53 | AM | 83% | 84% | $0.240 \pm 0.196$ | 55% | $0.477 \pm 0.653$ |
| | | FM | 62% | 100% | $0.101 \pm 0.052$ | 70% | $0.174 \pm 0.164$ |
| Bilateral matched valleys | 66 | AM | 42% | 96% | $0.136 \pm 0.099$ | 71% | $0.161 \pm 0.232$ |
| | | FM | 18% | 100% | $0.043 \pm 0.015$ | 67% | $0.399 \pm 0.419$ |

While the majority of significant AM correlation strengths were positive, a few bilateral matched pairs had negative correlation strengths (Figure 5A). Pair category had a significant effect on AM correlation strength, which was greatest in bilateral matched peaks, followed by bilateral matched valleys, and then ipsilateral adjacent peaks (one-way ANOVA, *F*_(2,103)_ = 18.56, *p* < 0.001; Tukey’s post-hoc tests returned *p* ≤ 0.041 for all comparisons). Significant FM correlation strengths were always positive (Figure 5B); that is, as the instantaneous frequency of a peak or valley decreased or increased, the same occurred in the other ear. FM correlation strength was significantly greater for bilateral matched peaks than valleys (unpaired *t*-test assuming equal variance, *t*_(42.0)_ = 5.71, *p* < 0.001). The strongest AM and FM correlations for bilateral matched peaks occurred near the mean SOAE peak center frequency of 2.5 kHz.

**Figure 5:**
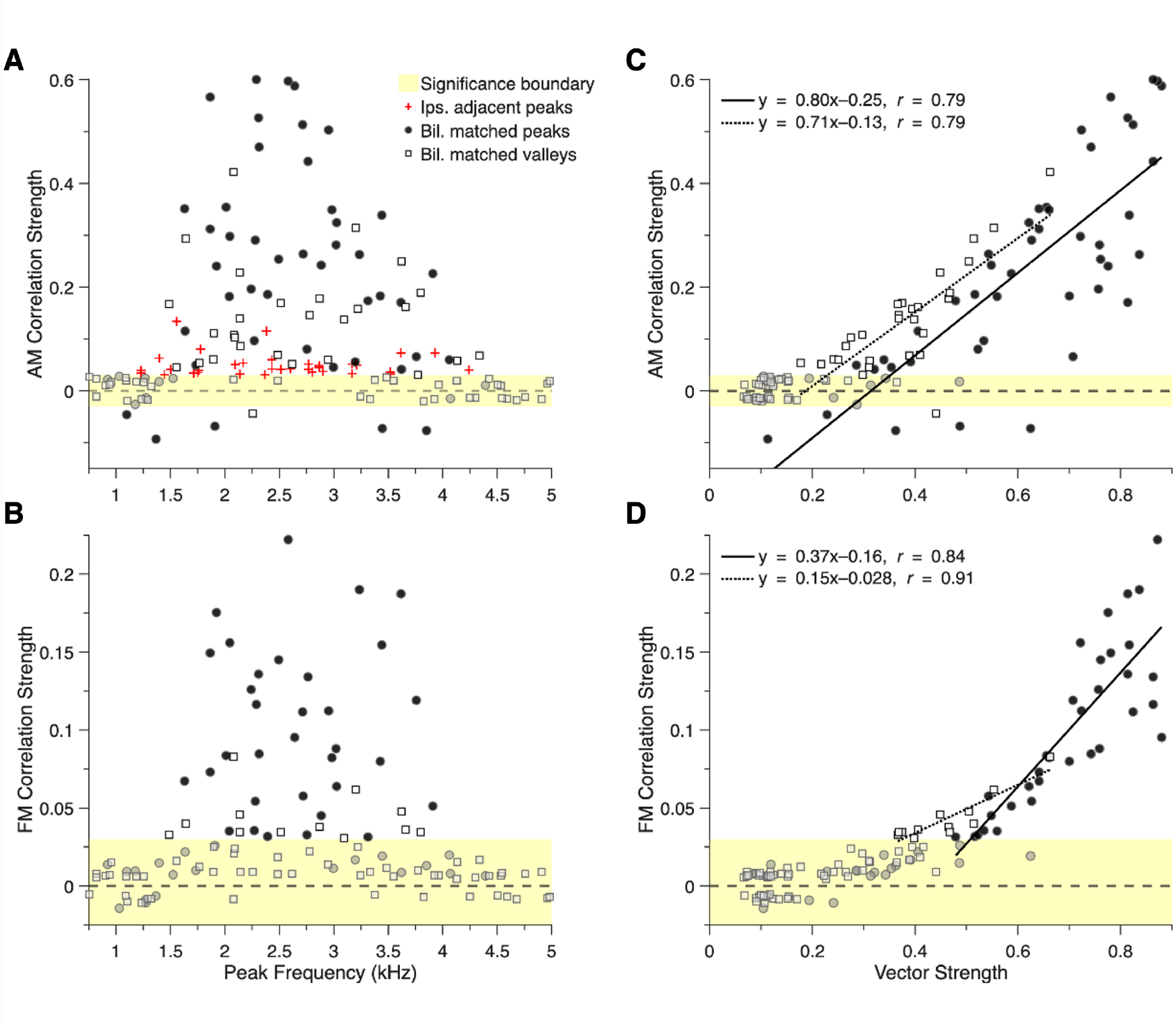
SOAE peak envelope (AM) and instantaneous frequency (FM) fluctuation correlation strengths computed from bilaterally measured waveforms exhibited relationships with peak frequency and vector strength. In all panels, horizontal dashed lines at 0 designate the boundary between positive and negative correlations and sub-threshold correlation strengths (values < ±0.03) are faded and highlighted with yellow shading. Significant and sub-threshold correlation strengths are shown for bilateral matched peaks (solid circles) and valleys (open squares). Due to the majority of ipsilateral adjacent peak pairs not meeting the significance criterion (see Table 1), only significant values are shown (red crosses). (**A– B**) Significant AM fluctuation correlations (**A**) occurred in all pair categories and could be positive or negative for bilateral matched pairs, while FM fluctuation correlations (**B**) only occurred in bilateral matched pairs and were always positive. The greatest correlation strengths for bilateral matched peaks occurred near the mean peak centre frequency of 2.5 kHz. Note that for AM correlations between ipsilateral adjacent peaks, values are plotted as the mean center frequency of the pair. (**C–D**) Vector strength was strongly and positively correlated with AM (**C**) and FM (**D**) fluctuation correlation strength for both bilateral matched peaks and valleys. Pearson correlation coefficients and linear regression fits for bilateral matched peaks (solid lines) and valleys (dotted lines) were computed only for fluctuation correlation strengths ≥ ±0.03. Linear correlations were statistically significant for the four relationships shown (all *p* < 0.001).

Consistent with our observations of a similar pattern for vector strengths, which may be driven by a relationship with peak SNR, AM and FM correlation strengths were positively related to peak SNR (AM: *r*_(42)_ = 0.38, *p* = 0.012, FM: *r*_(31)_ = 0.58, *p* < 0.001). Higher fluctuation correlation strengths were also associated with greater vector strengths in bilateral matched pairs. As shown in Figure 5C, there were strong positive relationships between AM fluctuation correlation strength and vector strength for bilateral matched peaks and valleys (both *r* = 0.79, *p* < 0.001). These relationships were even stronger for FM fluctuation correlations (bilateral matched peaks: *r*_(31)_ = 0.84, *p* < 0.001, bilateral matched valleys: *r*_(10)_ = 0.91, *p* < 0.001; Figure 5D). Such is likely a consequence of both FM fluctuation correlations and vector strengths deriving from the phase relationships between bilaterally measured SOAE waveforms.

Fluctuation analyses provided the correlation function lags (*τ*), which allowed us to examine the relative delays between pairs of filtered peaks or valleys. Most ipsilateral adjacent peak pairs had positive AM correlation delays (Table 1), such that the lower-frequency peak led the higher one, but delay direction was variable for both AM and FM correlations in bilateral matched pairs. AM correlation delays were longer than FM correlation delays for bilateral matched peaks (Table 1). Furthermore, we observed that greater correlation strengths were associated with shorter delays between waveforms (marker size represents correlation strength in Figure 6A and B).

**Figure 6:**
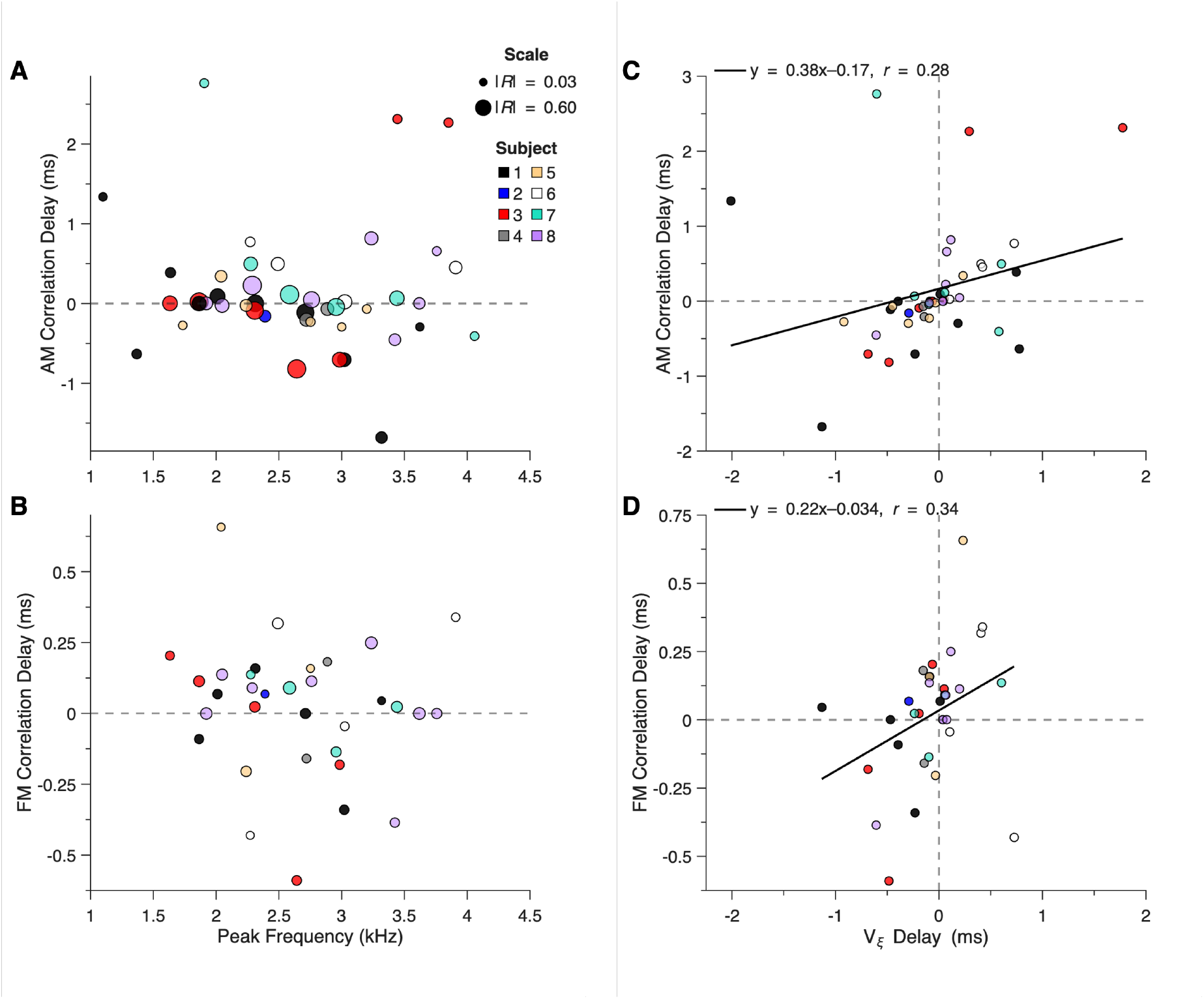
SOAE peak envelope (AM) and instantaneous frequency (FM) correlation delays between bilateral matched peaks were frequency-independent but related to the time-delayed vector strength, *V*_*ξ*_. Fluctuation correlation and *V*_*ξ*_ delays were computed as the left ear relative to the right ear, with positive delays (> 0 ms, see dashed lines) representing a peak from the left ear leading a peak from the right ear. (**A–B**) Fluctuation correlation delays plotted as a function of frequency for AM (**A**) and FM (**B**) correlations. Marker size is scaled according to correlation strength (|*R*|), which was inversely related to the delay such that stronger correlations had shorter delays. Each subject is represented by a unique color (see legend), illustrating that delays could be positive or negative for a given individual at different frequency bands. (**C–D**) *V*_*ξ*_ delays were weakly associated with AM (**C**) and FM (**D**) fluctuation correlation delays. Pearson correlation coefficients and linear regression fits between significant fluctuation correlation delays (correlation strength ≥ ±0.03) and *V*_*ξ*_ delays were computed for bilateral matched peaks (solid lines). Linear correlations were not significant (both *p* > 0.050).

Similar to the *V*_*ξ*_ delays (Figure 4B), correlation delays for bilateral matched peaks were not linearly related to peak frequency and could switch directions at adjacent frequency bands (Figure 6A and B). While frequency-independence might be expected if one ear dominantly drove the other, our results were not consistent with this; we found that the delays between bilateral matched peaks were not uniform across frequency for most subjects, with the majority exhibiting positive and negative delays for AM (five subjects, Figure 6A) and FM correlations (seven subjects, Figure 6B). When we compared correlation delays to *V*_*ξ*_ delays for bilateral matched peaks, we observed weak positive relationships (see labelled trend lines in Figure 6C and D, *p* > 0.05 for both). Some peaks exhibited correlation delays with opposite signs relative to the *V*_*ξ*_ delays. In Figure 6C, there were a handful of outliers where AM correlation delays exceeded ±1 ms. These values were associated with AM correlations that were classified as negative or indeterminate, reflecting more complex relationships between waveforms. Although all FM correlations were classified as positive, we still observed cases where the direction of the correlation delay was not consistent with the direction of the *V*_*ξ*_ delay (Figure 6D).

## 4. Discussion

### 4.1. Synchronisation in the Anole Auditory Periphery

The ears of green anole lizards exhibit binaural synchronization between their SOAEs. The coincidence of high vector strengths and strong fluctuation correlations with bilateral matched peaks was indicative that binaural coupling could manifest as frequency-dependent phase-locking between the two ears. Furthermore, binaural synchronization did not simply arise because one ear dominantly drove the other. Rather, there could be multi-band crosstalk between the two inner ears. Such results support that binaural synchronization arises due to the IAC providing a pathway for frequency-dependent interactions between the ears.

We interpret these results as indicative that the SOAE spectrum measured at one ear could be the product of energy from both ears. When evaluating fluctuation correlations, we consistently found that relationships across ears were stronger than those within a given ear. Given that SOAE generators appear to readily entrain to external stimuli (e.g., Köppl and Manley, 1994), activity in the contralateral ear could likewise serve as a source of entrainment. That is, not only do the generators in one ear adjust their characteristic frequencies to synchronize with their neighbours (e.g., Wit and Bell, 2023), but they may also adjust their frequencies to synchronize with those in the contralateral inner ear. While many models of SOAE generation emphasize coupling between nearby elements in local-oscillator frameworks (Gelfand et al., 2010; Talmadge et al., 1991; Vilfan and Duke, 2008; Wit, 1990; Wit and van Dijk, 2012), we consider that binaural synchronization may serve as a form of “global” coupling (Shera, 2022).

We propose an expanded framework for SOAE generation in the green anole lizard that accounts for contributions from two active inner ears. As illustrated in Figure 7, this model incorporates multiple sources of motion (panel A) and levels of coupling that could give rise to synchronization under the appropriate conditions. Local coupling between oscillating hair cells (bundle motion, *d*_*B*_) with similar characteristic frequencies can lead to synchronized groups that emit at a common frequency (Figure 7B; Talmadge et al., 1991; Vilfan and Duke, 2008). Importantly, rather than restricting “local” to mean an individual, oscillating hair cell, we extend our definition to include the coupling between neighboring hair cells as well as their surrounding environment (Shera, 2022), encompassing other cellular elements that can contribute cooperatively to active force generation within an ear (Bergevin et al., 2025).

**Figure 7:**
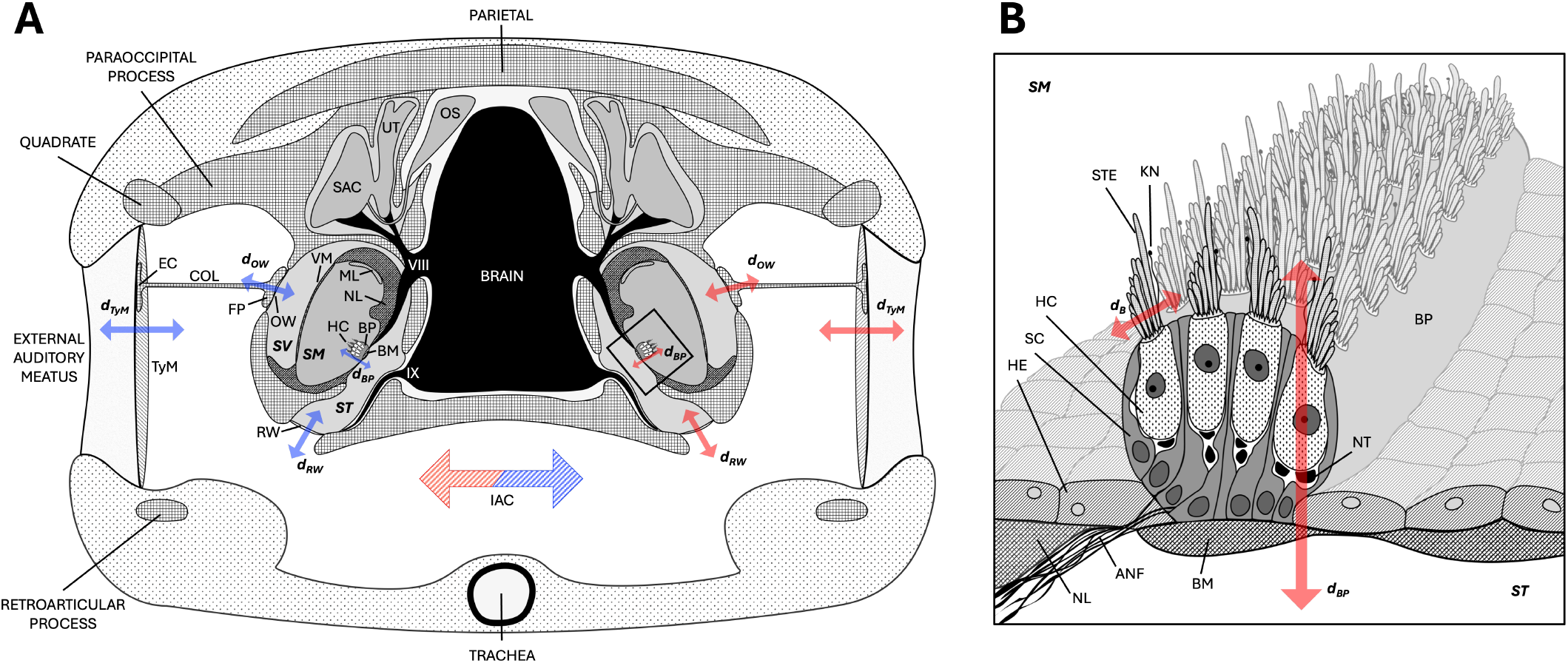
Schematic of the auditory system of the green anole lizard illustrating axes of motion and pathways for coupling. (**A**) Section of the head showing the relation between the external auditory meatus and inner ear on each side and the connection between the two ears via the IAC. Bidirectional arrows identify sources of motion that contribute to the transmission of acoustic energy in each ear (left in blue, right in red) as well as the ability for acoustic energy from one ear to affect the other by traversing the IAC (bicolour striped arrow). (**B**) Enlarged illustration of the portion of the right inner ear identified by the black box in panel A with enhanced anatomical detail. Note the absence of an overlying tectorial membrane on the hair cell bundles and their bidirectional orientation. **VII**: 8th nerve (vestibulo-cochlear nerve); **XI**: 9th nerve (pharyngeal nerve); **ANF**: auditory nerve fiber; **B**: hair cell bundle, *d*_*B*_: radial motion of the bundle (Aranyosi and Freeman, 2005); **BM**: basilar membrane; **BP**: basilar papilla, *d*_*BP*_: transverse motion of the basilar papilla (Aranyosi and Freeman, 2005); **COL**: columella; **EC**: extracolumella; **EC**: hyaline epithelial cell; **FP**: columella footplate; **HC**: hair cell; **KN**: kinocilium; **IAC**: interaural cavity; **ML**: macular lagena; **NL**: neural limbus; **NT**: nerve terminal; **OS**: otic sac (vestibular system); **OW**: oval window, *d*_*OW*_ : displacement of the oval window; **RW**: round window, *d*_*RW*_ : displacement of the round window; **SAC**: sacculus (vestibular system); **SC**: supporting cell; **SM**: scala media (endolymph); **ST**: scala tympani (perilymph); **SV**: scala vestibuli (perilymph); **STE**: stereocilium; **TyM**: tympanic membrane, *d*_*TyM*_: displacement of the tympanic membrane; **UT**: utricle (vestibular system); **VM**: vestibular membrane. Anatomical illustrations for panels A and B adapted from Baird (1970) and Weiss et al. (1974), respectively. Not to scale.

Each group of synchronized hair cells represents an oscillatory unit. A unit could couple not only to other groups within the same inner ear, possibly giving rise to ipsilateral fluctuation correlations between adjacent SOAE peaks, but also to those at similar frequencies in the contralateral inner ear, resulting in vector strength peaks and bilateral fluctuation correlations. Within a given ear, inter-group coupling is presumed to depend on hair cells being tuned to similar characteristic frequencies. Due to tonotopy, these cells are arranged in proximity to each other and are subject to a similar noisy environment (e.g., the Brownian motion of hair cell bundles due to thermal fluid agitation; Altoè and Shera, 2024; Bergevin et al., 2025). Possible coupling sources include the basilar membrane or the local endolymphatic fluid, which contribute to basilar papilla motion in a given ear (*d*_*BP*_, Figure 7B; Bergevin and Shera, 2010; Fruth et al., 2014; Wit and Bell, 2022). While hair cells in the frequency region corresponding to SOAE generation in anole lizards lack an overlying tectorial membrane (Manley, 1997), this could serve as another coupling source in other species (van Dijk and Manley, 2013).

Although hair cell groups at similar frequencies in both ears are spatially separated and subject to noise in their respective local environments, we observed synchronization between many bilateral matched peaks and valleys. Such relationships may be the consequence of frequency matching (Pikovsky et al., 2001), resulting in the coupling between distant but like elements being stronger than that between adjacent but unlike elements. If this is the case, ipsilateral fluctuation correlations would be weaker than bilateral fluctuation correlations, as we observed in the green anole. The rarity of FM fluctuation correlations between ipsilateral adjacent peaks (none observed here; three in Bergevin et al., 2025) relative to bilateral matched peaks and valleys suggests that frequency matching plays an important role in these relationships.

Binaural synchronization most likely occurs because the two ears are weakly coupled via the IAC (Bergevin et al., 2020; Roongthumskul et al., 2019). While other sources, such as bone conduction, could contribute, past modelling indicates that the motion of one tympanic membrane (TyM; *d*_*TyM*_) is sufficient to drive the other (Figure 7A; Bergevin et al., 2020). We assume this is the dominant pathway for binaural coupling despite our report of attenuation due to interaural sound transmission (approximately 10—25 dB attenuation between 0.9—4.4 kHz when measured at the external auditory meatus; Figure A.8).

The presumed pathway for sound energy to exit the inner ear is via the oval window (*d*_*OW*_ ), which drives the columella and subsequently the TyM (*d*_*TyM*_), such that an emission is measurable at the external auditory meatus (Figure 7A). However, the round window could serve as an additional pathway for sound energy (*d*_*RW*_ ), entering the IAC. In lizards, the structure has been previously identified as the “secondary tympanic membrane” in recognition that it is not directly homologous to the mammalian round window (Baird, 1970). Energy from the round window could stimulate the ipsilateral TyM and possibly the contralateral TyM due to the coupling effects of the IAC (Figure 7A). The interplay between the two TyMs could create a feedback loop for sound energy within the IAC, which could underlie the large SOAE activity observed in lizards like the green anole (e.g., Manley, 2006).

The role of the round window in SOAE transmission should be evaluated in future experiments in green anole lizards. Although severing the columella via ossicular interruption can eliminate evoked emissions at the corresponding external auditory meatus in bobtail lizards (*Tiliqua rugosa*; Manley et al., 1993), it is unknown whether this would likewise eliminate the transmission of sound to the contralateral inner ear and, ultimately, the TyM. The persistence of binaural synchronization, as evidenced through vector strength and fluctuation correlations, after severing the columella in one ear, would demonstrate that the round window provides another pathway for interaural sound transmission.

### 4.2. Variations in Synchronization

While we found evidence of binaural synchronization in each ear studied, the extent to which it was present was variable. Some anole ears were far more “synchronized” than others (e.g., compare the vector strengths of subjects 1 and 2 in Figure SI A.9A and B, respectively). Furthermore, binaural synchronization did not explicitly depend on spectral similarity; we observed elevated vector strength and significant fluctuation correlations when comparing dissimilar regions (e.g., high vector strength between 3–4 kHz for subject 6 in Figure A.9F). Such variability could account for earlier reports that SOAEs from bobtail lizards were generated solely by the ipsilateral ear without any contributions due to crosstalk from the contralateral ear’s emissions (Köppl and Manley, 1993). This conclusion was based on the observation that abolishing SOAEs in one ear did not produce a consistent difference in the contralateral spectra measured before, during, or after the abolishment. However, Roongthumskul et al. (2019) demonstrated that manipulating the ear canal pressure at one ear to suppress or attenuate its emissions could have varying effects on the contralateral SOAE spectrum. With this in mind, the “null” results from bobtail lizards lend further support to our findings that binaural synchronization occurs in frequency-dependent channels and can be variable across individuals.

It was proposed by Roongthumskul et al. (2019) that binaural synchronization of SOAEs depended on the strength of each peak and how susceptible the generators were to drift from their characteristic frequency to synchronize with another (i.e., detuning; Pikovsky et al., 2001). SOAEs from green anole lizards were of comparable magnitude to those from the tokay gecko, and, in support of predictions made by Roongthumskul et al. (2019), we found weak positive correlations between our proxy for peak strength (SNR) and vector strength as well as fluctuation correlation strength. However, maximum vector strengths were found to be higher in the green anole than in tokay geckos, despite the anole’s smaller and relatively simpler basilar papilla (Wever, 1978). This could be explained by the generators in the green anole being more easily detuned from their characteristic frequencies by a bilateral counterpart than those in the tokay gecko.

In the anole, fewer hair cells contribute to each emitted peak, and their bundles are relatively large and densely packed compared to the gecko (Baird, 1970; Negandhi et al., 2018). The proximity of neighboring bundles in the anole could increase the probability of hair cells detuning each other, synchronizing to emit at a common frequency. Furthermore, unlike the tokay gecko, the green anole does not have an overlying tectorial membrane on its hair cell bundles (Manley, 1997). The absence of a tectorial membrane has been previously proposed to weaken reactive coupling and perhaps increase resistive coupling between hair cells, producing more SOAE peaks with narrower inter-peak spacing (Wit et al., 2020). Hair cells in the anole could be more likely to entrain to an emission at a nearby frequency in the contralateral ear, resulting in greater synchronization between bilateral matched peaks. Conversely, the salletal structures overlying hair cell bundles in the gecko are thought to strengthen the coupling between adjacent hair cells (Chiappe et al., 2007; Köppl and Manley, 1993; Wit et al., 2020), which would provide greater opposition to detuning due to binaural coupling.

### 4.3. Not all Bilateral Matched Peaks are Binaural

We interpreted the properties of many bilateral matched peaks as indicative that they occurred due to the oscillators in each ear mutually adjusted their characteristic frequencies to emit at a common frequency, or even arose from binaural synchronization itself as seen in geckos (Roongthumskul et al., 2019). In most cases, both ears exhibited similar *C*_*ξ*_ values at the same frequencies, time-shifted vector strengths (*V*_*ξ*_) were highest near 0 ms, FM correlation strengths were positive, and fluctuation correlation delays were near 0 ms. However, some bilateral matched peaks exhibited properties suggestive that one ear entrained the other at those frequency bands. Some *V*_*ξ*_ and fluctuation correlation delays substantially deviated from 0 ms, and, contrary to expectations, the strongest *C*_*ξ*_ values for a given subject did not necessarily occur at the same frequencies as the highest vector strengths, and some ears exhibited distinct *C*_*ξ*_ values at the same frequencies. While many SOAE peaks exhibit phase stability over time, the *C*_*ξ*_ in one ear could be independent of the *C*_*ξ*_ in the other ear. Such could reflect the underlying “source” of a bilateral matched peak pair being one inner ear, which then elicits a response in the other.

Occasionally, a bilateral matched peak pair did not have a corresponding peak in the vector strength. In three illustrative cases where this occurred, the SOAE peak amplitude was higher in one ear, and there was a clear peak in that ear’s *C*_*ξ*_, but not for the contralateral ear (e.g., see yellow arrows for subjects 1, 4, and 5 in the second row of panels in Figure A.9A, D, and E). Furthermore, these three peak pairs had negative AM fluctuation correlations (although the pairs from subjects 4 and 5 did not meet criterion). However, further work is needed to understand how such correlations arise. It is unlikely that energy was passively leaking from one inner ear to the contralateral TyM without driving synchronization between ears because most SOAE peaks were too small to generate measurable counterparts in the contralateral ear (due to peak amplitudes lower than the attenuation reported in Figure A.8). Nevertheless, these cases suggest that some bilateral matched peaks were the result of the ears coincidentally emitting at the same frequencies and were not attributable to binaural synchronization.

### 4.4. Synchronization Beyond the Peaks

Binaural synchronization was not restricted to the SOAE peaks themselves. Although weaker than for bilateral matched peaks, vector strengths and fluctuation correlations could be high at the valleys between peaks (see square markers in Figure 5C and D). Consistent with previous work in anoles (Bergevin et al., 2025), bilateral fluctuation correlations were only apparent when the valley between peaks was above the presumed microphone noise floor (e.g., see the distance between the grey dashed lines and the measured SOAEs in Figure 1B). Furthermore, fluctuation correlations did not occur when the valleys were at frequencies above or below the maximum or minimum SOAE peaks, respectively.

The observed SOAE valleys likely constitute “baseline” emission activity (e.g., Köppl and Manley, 1993; Manley et al., 1996; Manley and Gallo, 1997), such that there is a broader underlying emission upon which the SOAE peaks are superimposed. While this baseline emission has been attributed to the presence of the tectorial membrane sallets in geckos (e.g., Köppl and Manley, 1993; Manley et al., 1996; Manley and Gallo, 1997), the absence of a tectorial membrane in the anole (Manley, 1997) suggests another source. One possibility is that baseline emissions arise due to weakly synchronized activity between the ears, creating a broad emission centered at the resonant frequency of the IAC (Roongthumskul et al., 2019; van Dijk and Manley, 2013).

In the gecko, Roongthumskul et al. (2019) predicted that the fundamental frequency of the IAC was between 3–4 kHz because phase differences for most bilateral matched peaks below 3 kHz were approximately 0.5 cycles and 0 cycles, or equivalently 1 cycle, at higher frequencies. In green anole lizards, phase differences were near 0.5 cycles between approximately 1.5–3.5 kHz, the frequency range where supra-threshold vector strengths were most prominent (bottom rows of Figure A.9). Phase differences near 0 or 1 cycle were poorly defined and associated with sub-threshold vector strengths. The persistence of anti-phase relationships between ears in anoles but not in geckos could be associated with the bidirectional polarity of hair cell bundles in anoles and the absence of a tectorial membrane, as shown in Figure 7B. If baseline emissions are attributable to binaural coupling, manipulating IAC coupling by occluding one ear or selectively suppressing its activity would not only disrupt binaural synchronization but also eliminate baseline emission activity.

### 4.5. Binaural Synchronization and Sound Localization

It has been reported that lizards have better sound-localization acuity than would be predicted from their head size and hearing range (e.g., Christensen-Dalsgaard et al., 2021), and this can be attributed to binaural coupling via the IAC. Multiple models of sound localization predict that the directionality of their ears stems from the acoustic interactions of the two TyM due to binaural coupling (Christensen-Dalsgaard and Manley, 2008; Livens et al., 2019; Vedurmudi et al., 2016; Vossen et al., 2010). However, this is primarily considered a passive process, with models elaborating on Fletcher and Thwaites (1979)’s pressure gradient receiver. Few models consider how the mechanisms that underlie active hearing in one ear, let alone two, could contribute.

Binaural synchronization would imply that the active mechanism in one ear can interact with the other, which has important implications for low-level sounds where the effects of the active mechanism (i.e., enhanced sensitivity and frequency selectivity) are most pronounced (Hudspeth, 2008). In their mathematical model of the gecko’s auditory periphery, Roongthumskul et al. (2019) demonstrated that binaural sound localization derived from contributions from two ears; sensitivity to weak sounds and determination of sound source were dependent on the activity of both. A greater degree of binaural synchronization at a given frequency band might result in stronger directionality. Our finding that binaural synchronization could be asymmetrical at a given frequency band (i.e., driven by one ear entraining the other) might result in the dominant ear contributing more to sound-source localisation. In cases where SOAE peaks appear to arise explicitly because of binaural synchronization, interaural cues may be dependent on the presence of IAC coupling. Thus, active contributions from each inner ear may be an important factor in the strong directional sensitivity observed in lizards.

## 5. Conclusions

Binaural synchronization is at work in the auditory system of the green anole lizard. The observed relationships between vector strength and fluctuation correlation strength supported that the two ears have frequency-dependent interactions. Furthermore, our results indicated that binaural coupling was stronger than that between generators within a given ear, suggesting stronger synchronization between SOAE-generative elements across ears compared to the generators within an individual ear. We propose a model for binaural SOAE generation in the green anole that accounts for multiple levels of coupling in the auditory periphery. While cellular cooperativity can shape the interactions between adjacent hair cells, these cells are also subject to binaural synchronization. Consequently, emissions measured at one ear can reflect not only the activity of that ear but also the contralateral ear.

## Appendix A. Supplementary Figures

Additional Figures A.8 and A.9.

**Figure A.8:**
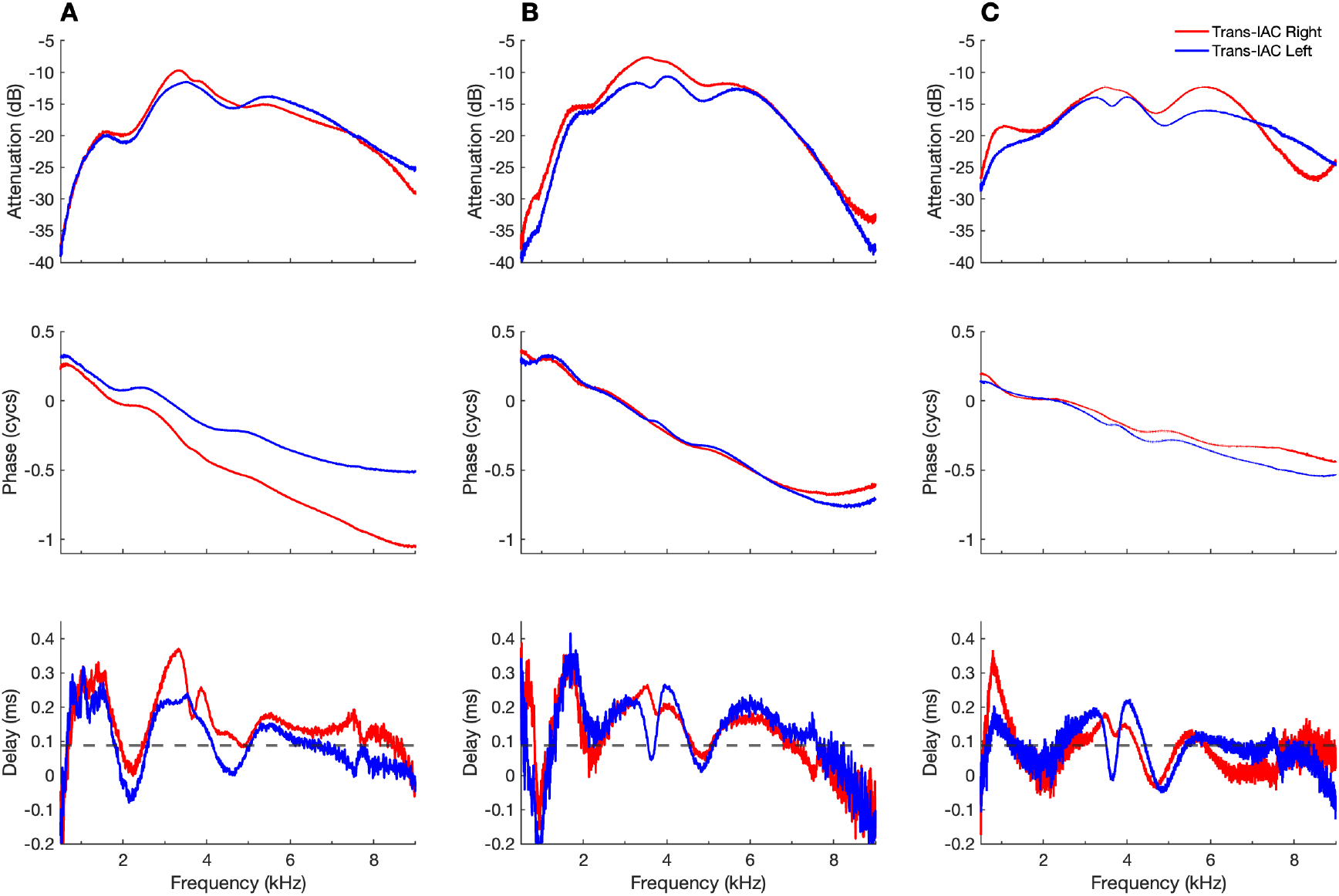
Attenuating effects of the interaural cavity (IAC) in three representative green anole lizards. Panels A–C correspond to Subjects 4, 5, and 8 in Figure A.9D, E, and H, respectively. (**Top row**) Acoustic attenuation was frequency-dependent, being lowest approximately 2–4 kHz, but exhibited subject-specific variation. (**Middle row**) Phase values decreased with increasing frequency. (**Bottom row**) Phase gradients were used to estimate interaural sound transmission delays across frequencies. Dashed lines in these panels represent delay of 0.090 ms, the estimated sound propagation delay based on the anole’s head size.

**Figure A.9:**
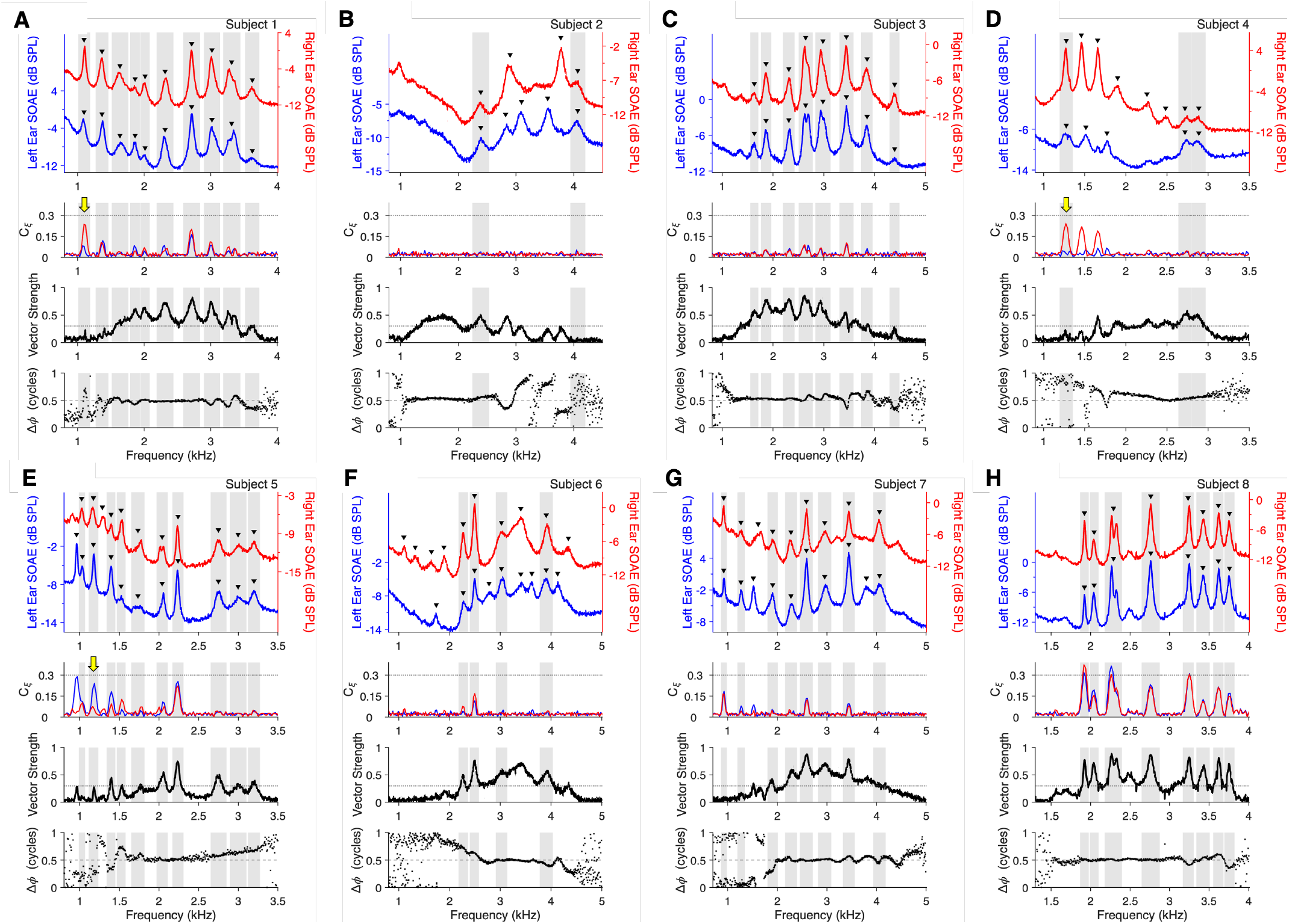
Overview of binaural SOAE properties in green anole lizards (*N* = 8). Subject ID for each individual is stated above the top panels. Subject 1 is the same lizard as shown in Figure 3. **(Top row for each subject)** Averaged SOAE spectra measured simultaneously at the left (blue; left vertical axes) and right (red; right vertical axes) ears. Note that the scales for the vertical axes were offset to avoid visual overlap. SOAE peaks are identified by black triangles above each waveform. When SOAE peaks were filtered at the same frequencies in both ears, they were classified as bilateral matched peaks and grey shading indicates their bandwidths across all panels. **(Second row)** Time-delayed phase coherence (*C*_*ξ*_) for the SOAE waveforms from the left (blue) and right (red) ears. Yellow arrows are included in some panels to emphasize cases where the *C*_*ξ*_ value at a bilateral matched peak was higher in one ear than the other, as discussed in Section 3.2. *C*_*ξ*_ was computed with a frequency resolution of 21.5 Hz, while all other data illustrated here were computed with a resolution of 5.4 Hz. **(Third row)** Vector strength between the SOAE waveforms (black) across frequencies. The dashed line at 0.3 represents the threshold for identifying vector strength peaks. **(Fourth row)** The phase differences (Δ*ϕ*) between SOAE waveforms were corrected to a range of 0–1 cycles (black dots), with a dashed grey line to demarcate 0.5 cycles.

## Appendix B. Acknowledgements

Supported by the Natural Sciences and Engineering Research Council of Canada (NSERC) to REW and CB.

